# Wnt stimulation and inhibition in the development and phenotype of patient-derived gallbladder organoids

**DOI:** 10.64898/2026.04.06.716840

**Authors:** Ankita Dutta, Payel Guha, Akshaya Vijayan Selvarajan, Nandita Chowdhury, Pritha Banerjee, Shrabanti Sarkar Ghosh, Archana Kumari Shaw, Debdutta Ganguli, Uma Sunderam, Manas Kumar Roy, Sudeep Banerjee, Rajgopal Srinivasan, Paromita Roy, Vaskar Saha, Anindita Dutta, Dwijit GuhaSarkar

## Abstract

Gallbladder cancer (GBC) is a highly lethal malignancy with limited experimental models to study disease biology or evaluate therapeutic responses. Although canonical Wnt activation is commonly used for patient-derived organoid (PDO) development and expansion, gallbladder PDOs has also been generated under Wnt-inhibitory conditions. No comparative assessment has determined how Wnt pathway modulation influences gallbladder PDO development, phenotype or drug response. This study systematically compared the impact of canonical Wnt activation (WNT^Act^ medium containing CHIR99021) versus inhibition (WNT^Inh^ medium containing DKK1) on the establishment, propagation, molecular features and therapeutic responses of PDOs generated from malignant or non-malignant gallbladder tissues derived from the same patient.

Both media supported successful PDO generation with comparable efficiency, preserving biliary epithelial functions and marker expression. Transcriptomic profiling confirmed selective enrichment of canonical Wnt target genes in PDOs generated in WNT^Act^ cultures. WNT^Act^ conditions enabled markedly superior long-term propagation, whereas WNT^Inh^ cultures more consistently retained the dysplastic features in malignant samples. Gemcitabine response assays demonstrated significantly greater drug sensitivity in PDOs grown in WNT^Act^ medium, a phenotype reversible upon media switching but requiring extended adaptation, indicating a dynamic and context-dependent influence of Wnt signaling on chemotherapeutic vulnerability. Collectively, the findings reveal a trade-off between long-term propagation and histological fidelity in gallbladder PDOs and show that Wnt signaling modulates gemcitabine sensitivity in a reversible manner. This comparative framework provides practical guidance for selecting culture conditions for gallbladder PDO based disease modelling and precision oncology applications.

## Introduction

Gallbladder cholangiocyte organoids (GCOs) and gallbladder carcinoma organoids (GBCOs) are valuable model systems for investigating gallbladder cancer pathogenesis (1), performing drug response testing (2), identifying novel biomarkers and developing alternative therapeutics to improve the patient outcomes. Generation of models that faithfully recapitulate the histopathological, functional, and molecular characteristics of the source tissue is reliant on appropriate signaling cues in the culture environment (3). Among these cues, Wnt signaling plays a critical role in organ development and tissue homeostasis (4). Under ex vivo conditions, Wnt signaling can be modulated through the addition of exogenous molecules (5). Conventionally, GCOs and GBCOs have been developed by activating the canonical Wnt signaling pathway (6,7). The efficiency of GBCO generation using this method is low (7). Patient-derived organoids (PDOs) have also been generated by inhibiting the canonical Wnt pathway but limited to GCOs (8,9). Wnt activation with the GSK3 inhibitor CHIR99021 potentially maintains cell stem/progenitor property, promotes self-renewal and improves long term passaging capability but suppresses terminal differentiation (6). In contrast Dickkopf-1 (DKK1), a Wnt antagonist induces cellular differentiation, maintains structural fidelity but may decrease self-renewal capacity (8). Because gallbladder tissue biology is less characterized than intestine or liver, testing both conditions (activation vs inhibition) is necessary to determine the right culture conditions to generate optimal models. In this study, we performed a comparative analysis of two organoid growth media compositions: one containing CHIR99021 to activate canonical Wnt signaling (WNT^Act^) and another containing DKK1 to inhibit the pathway (WNT^Inh^). Our aim was to determine suitable culture conditions for the development of patient-derived GCO and GBCO models.

## Materials and methods

### 2.1 Patient identification and sample collection

Informed consent was obtained from patients undergoing surgical gallbladder removal at the Tata Medical Center Kolkata (TMCK) between January 2021 to January 2025. Ethical approval was obtained from Institutional Review Board (EC/TMC/65/16 and 2021/TMC/216/IRB3). Tissue samples were collected and transported in cold sterile tissue transport media as mentioned previously (10).

### 2.2 Organoids generation and expansion

Epithelial cells from the surgically resected gallbladder tissue were isolated and 20,000 – 50,000 cells/well were seeded in reduced growth factor Geltrex^TM^ (A1413202, Gibco) dome (20µL cell suspension in 40µL matrix) in a 24-well culture dish (3526, Corning) as previously described (10). Cultures were grown in two different types of organoid growth media (1ml/well) - WNT^Act^ containing GSK3 inhibitor CHIR99021 and WNT^Inh^ containing Wnt inhibitor DKK1 **[Supplemental Table S1]**, compositions of which were adapted from previous reports (2,9). PDO cultures were expanded and cryopreserved. Organoids were also generated from cryo-preserved primary cells as described before (10).

### 2.3 Size analysis

Microscopic images of the PDOs were captured from a 24-well culture dish in bright-field mode on Nikon Eclipse TS2, 10X air objective within 7 days of initial seeding. Diameters were measured in two perpendicular axes for each PDO using FIJI software and mean diameter was calculated. PDOs for measurement were chosen from similar regions of interest (ROI) for each paired culture with a minimum of 10 PDOs in each ROI. A minimum of 50µm mean diameter was considered as an eligibility criterion to be considered as a PDO in this analysis.

### 2.4 Growth analysis

PDOs were split at 1:2 ratio when the confluence reached 70 to 80%. Media change was given every 3 to 4 days. Organoid growth curve was plotted as number of passages reached against culture duration. Culture survival probability was analyzed by Kaplan-Meier survival analysis using GraphPad Prism 10. Degenerating cultures were counted as event and the growing or cryopreserved cultures were censored for the analysis.

### 2.5 Histological analysis and immunohistochemistry

Organoids were harvested and fixed in 10% Neutral Buffered Formalin (NBF) and proceeded for formaldehyde fixed paraffin embedded (FFPE) block preparation, microtome sectioning, and hematoxylin-eosin (HE) staining or immunohistochemistry (IHC) (10). For IHC antibodies were used against CK7 (1-CY163-07, Diagomics, 1:100), CK20 (MU315-UC, BioGenex, 1:50) and Ki67 (MIB-1, Z2305ML, Zeca corporation, 1:200). Ki67 proliferation index was calculated by hotspot method using online software tool IHCExpert (https://www.ihcexpert.com/).

### 2.6 Functional assays

Active membrane transport function of PDOs were determined by checking efflux of Rhodamine 123 (R8004, Sigma-Aldrich), a fluorescent substrate of p-glycoprotein (p-gp or ABCB1) transmembrane pump. Organoids were incubated with fresh medium containing 100μM Rhodamine 123 dye and fluorescent microscopy was performed to assess efflux. To confirm specificity, PDOs were incubated with 10μM verapamil hydrochloride (V4629, Sigma-Aldrich), an inhibitor of the pump to see if the efflux could be blocked. Alkaline phosphatase assay was performed using BCIP/NBT Color Development Substrate (5-bromo-4-chloro-3-indolyl phosphate/ nitro blue tetrazolium) (S3771, Promega) as described previously (10). Organoids were incubated with fresh colorless medium containing the substrates for 1 hour at 37°C. Images were captured in bright-field mode using Leica DMi8. Gamma glutamyl transferase activity was measured using the Gamma Glutamyl Transferase Assay Kit (colorimetric) following the manufacturer’s instructions (ab241029, Abcam) and as described previously (10).

### 2.7 qRT-PCR

RNA extraction, cDNA preparation and q-RTPCR was performed as previously described (10). Differential gene expression relative to the reference gene and fold change in the gene expression relative to WNT^Act^ growth condition was calculated. Paired t-test using GraphPad Prism 10 was performed to evaluate the statistical significance.

### 2.8 mRNA sequencing and analysis

cDNA libraries were prepared from 200ng of total and sequenced on Illumina NextSeq 550 system (paired-end, 150 cycles x 2) (10). Principal component analysis (PCA) clustering was done after VST transformation of the 1000 highly variable genes (10) **[Supplemental table S2, S3A, S3B, S4A and S4B]**. Gene set enrichment analysis (GSEA) was performed using a customized set of 18 genes **[Supplemental Table S5]** on the size factor normalized counts matrix where genes with at least 10 counts in at least 25% of the samples were retained. Genes were ranked using the signal to noise metric, and statistical significance was assessed via phenotype-based permutations.

### 2.9 Drug response testing

8000 PDO-derived cells were seeded per well in 5% matrix in 96-well culture dish (Costar 3904). Viability was assessed after 72 hours of treatment with dimethyl sulfoxide (vehicle control) or gemcitabine (Sigma Aldrich-1003500321) at 7 different concentrations (10^-1^-10^5^ nM with 10-fold serial dilutions) using CyQUANT^TM^ Direct Cell Proliferation Assay (ThermoFisher Scientific-C35011). IC50, Hill slope, R_max_ (bottom) and R_min_ (top) were evaluated from the four-parameter logistic regression equation using GraphPad Prism 10. F-test was performed to evaluate the statistical significance.

## Results

### 3.1 Patient-derived gallbladder organoids were generated in both WNT^Act^ and WNT^Inh^ culture conditions

Patient-derived gallbladder organoids (PDOs) were successfully generated from surgically resected malignant and non-malignant tissues using both WNT^Act^ and WNT^Inh^ organoid culture media [**Supplemental Table S1**]. Tissue sources represented a spectrum of gallbladder pathologies, including chronic cholecystitis (CC), xanthogranulomatous cholecystitis (XGC), intracholecystic papillary tubular neoplasm (ICPN), adenocarcinoma (AdCa), adenosquamous carcinoma (AdSqCa), and histologically normal gallbladder. Organoid morphology often varied according to the pathology of the source tissue. Organoids derived from invasive carcinoma (AdCa) or pre-invasive neoplasms (ICPN) typically displayed irregular, optically dense, or mixed morphologies, whereas those derived from non-malignant tissues (CC, XGC) formed more regular and translucent cystic structures [**Figure 1A**]. Organoid generation success in either medium showed no association with any specific pathology [**Figure 1A, Supplemental Table S6**]. GCOs were also successfully generated from cryopreserved primary cells and expanded in both culture media [**Figure 1B**]. Although organoid morphology differed across patients or pathologies, no consistent media-dependent differences were observed between paired PDO cultures derived from the same patient but grown in WNT^Act^ or WNT^Inh^ media [**Figure 1A, B**]. Mean organoid diameters, measured during early passages (passages <4) were also comparable between PDOs cultured in the two media [**Figure 1C**].

**Figure 1:**
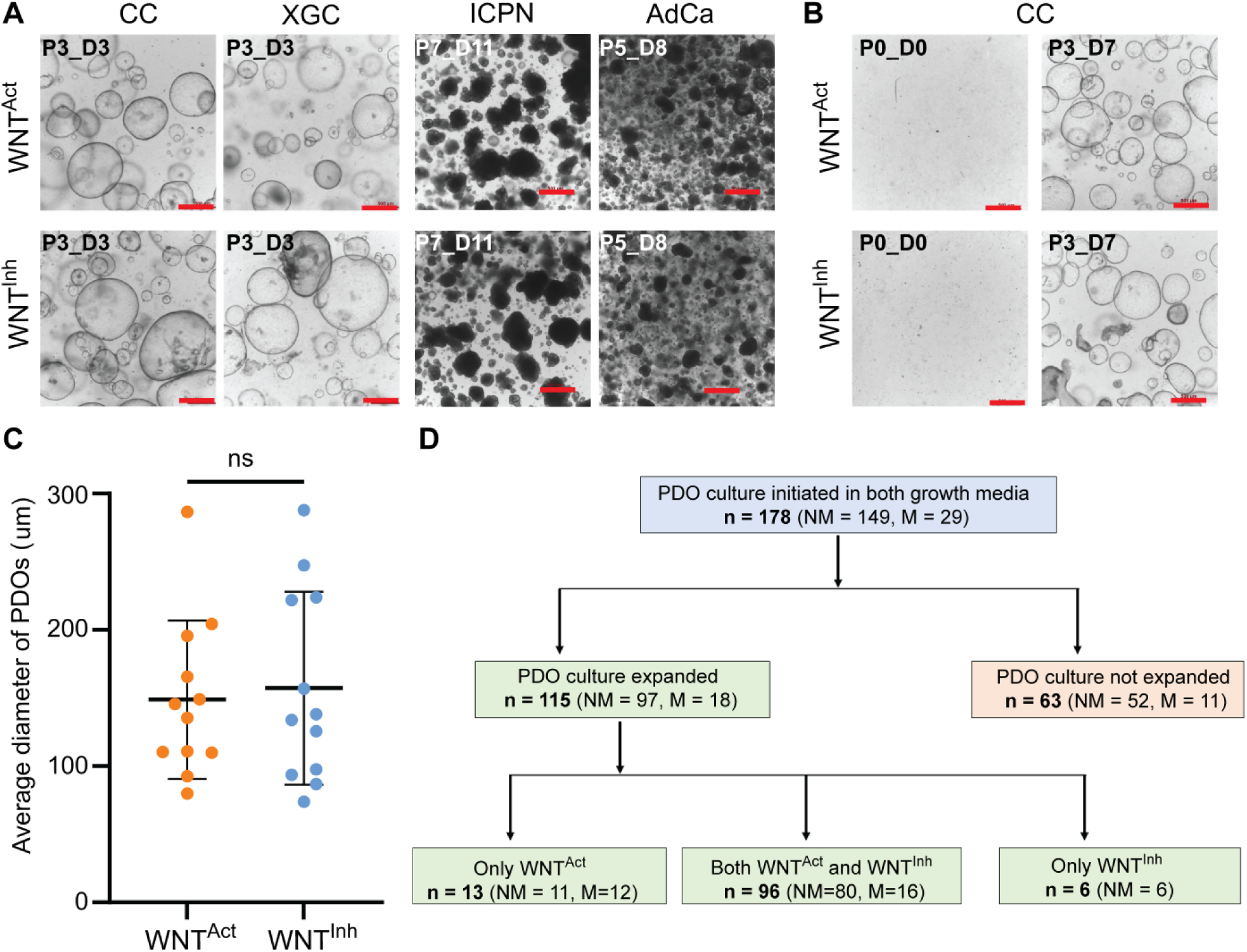
Establishment of patient-derived gallbladder organoids in WNT^Act^ and WNT^Inh^ media. Representative brightfield images of organoids derived from **(A)** freshly isolated cholangiocytes of CC, XGC, ICPN and AdCa gallbladder pathologies. Organoids from malignant tissues exhibit irregular morphologies, whereas those from non-malignant tissues form translucent cystic structures. But no consistent morphological difference was observed between the paired cultures grown in WNT^Act^ (top) and WNT^Inh^ (bottom). **(B)** Representative images of organoids derived from cryo-preserved primary cells of CC gallbladder pathology in WNT^Act^ (top panel) and WNT^Inh^ (bottom panel) growth media. **(C)** Dot plot showing quantification of mean organoid diameter in early passages (<P4) showing no significant difference between WNTAct and WNTInh cultures. Mean +/- SD (n =11). p-value ns (paired t-test). **(D)** Summary of organoid establishment and expansion success in both media, WNT^Act^ only, or WNT^Inh^ only. Scale bar: 500µm. Microscope – Nikon TS2, 4X objective. CC, chronic cholecystitis; AdCa, adenocarcinoma; XGC, Xanthogranulomatous cholecystitis; ICPN, intra-cholecystic papillary tubular neoplasm; P, passage number; D, days after last passage; ns, not significant; NM, non-malignant; M, malignant; PDOs, patient-derived organoids

Overall, 211 gallbladder PDO lines were generated from 115 patients. Organoid establishment was successful in at least one culture condition in 65% of patients (115/178). Among these, paired organoids were successfully generated in both in 83.47% (96/115) whereas 16.5% yielded organoids in only one condition (19/115; 13 in WNT^Act^ and 6 in WNT^Inh^) [**Figure 1D, Supplemental Table S6**]. For malignant gallbladder pathologies, GBCO generation was successful in at least one medium in 62% (18/29), of with 89% (16/18) were generated successfully in both the media. In total, 169 viable PDO lines (15 in WNT^Act^ alone, 1 in WNT^Inh^ alone, and 77 paired lines in both media) from 92 patients were successfully expanded and cryopreserved [**Supplemental Table S6**].

### 3.2 PDOs cultured in both WNT^Act^ and WNT^Inh^ media retain biliary epithelial functional characteristics and molecular marker expression

PDOs cultured in both WNT^Act^ and WNT^Inh^ media exhibited comparable functional characteristics of biliary epithelium. Luminal accumulation of Rhodamine-123, a known substrate of P-glycoprotein pump, indicated presence of functional P-glycoprotein pump activity in organoids grown under both conditions [**Figure 2A**]. Similarly, paired lines (n=3) expressed comparable levels of alkaline phosphatase [**Figure 2B**], and γ-glutamyl transferase activity [**Figure 2C**]. Gene expression analysis by quantitative RT-PCR confirmed consistent expression of key cholangiocyte markers (*KRT7, KRT19*, *CFTR*, and *SOX9*) and absence of hepatocyte markers (*ALB* and *AFP*) in PDOs grown in both culture conditions [**Figure 2D**]. Immunohistochemical analysis further confirmed expression of biliary epithelial markers CK7 and CK20 without detectable media-dependent variation [**Figure 2E**]. CK7 expression was uniformly strong across samples, whereas CK20 expression was patchy but consistent between paired lines derived from the same patient.

**Figure 2:**
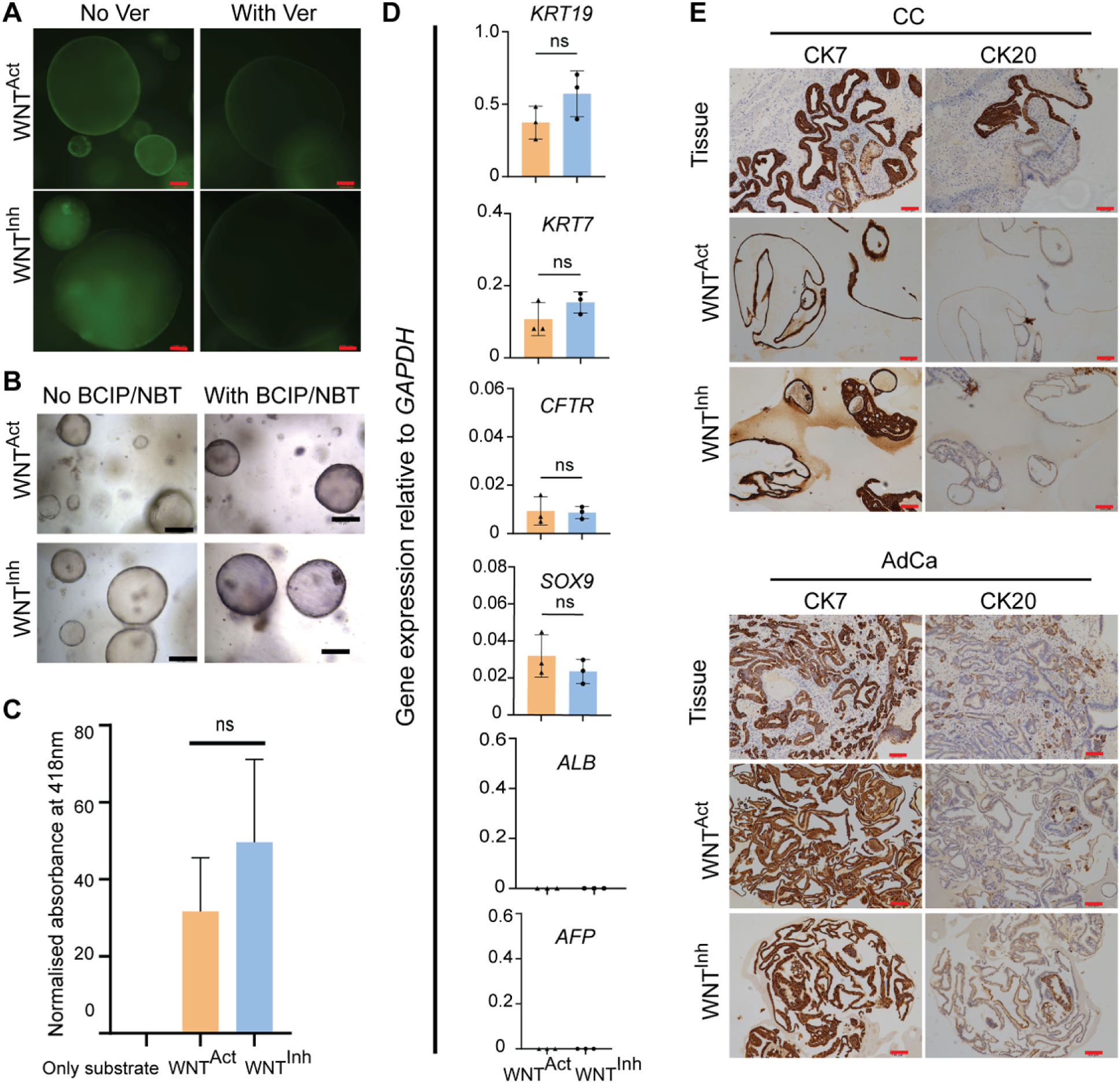
Functional and molecular characterization of PDOs cultured in WNT^Act^ and WNT^Inh^ media. Representative images showing **(A)** Rhodamine-123 - transport assay demonstrating active P-glycoprotein–mediated transport and luminal dye (green) accumulation in organoids cultured under both conditions **(B)** Assay using BCIP/NBT chromogenic substrate shows alkaline phosphatase activity in GCOs cultured in both WNT^Act^ and WNT^Inh^ media. **(C)** Bar plot depicting the GGT activity in paired GCOs developed in WNT^Act^ and WNT^Inh^ growth media. All the values have been normalized against the total cell count and plotted Mean +/-SD (n = 3); p-value ns (paired t-test). **(D)** qRT-PCR expression of cholangiocyte markers *KRT7, KRT19, CFTR, SOX9*, and hepatocyte markers *ALB* and *AFP*, plotted Mean +/- SD (n = 3); p-value ns (paired t-test) and **(E)** immunohistochemical staining confirming the expression of CK7 and CK20 in the PDOs developed in WNT^Act^ and WNT^Inh^ growth media as indicated along with their corresponding source tissue. The corresponding pathology is indicated above each panel. Scale bar - 200μm (black) and 100μm (red). Microscope – Nikon TS2, 10X objective (A); Leica DMi8, 10X objective (B and E). Ver, verapamil HCl; BCIP, 5-bromo-4-chloro-3-indolyl phosphate; NBT, nitro-blue tetrazolium; ALP, alkaline phosphatase; GGT, gamma glutamyl transferase; CC, chronic cholecystitis; AdCa, adenocarcinoma; GCOs, gallbladder cholangiocyte organoids; ns, not significant; PDOs, patient-derived organoids

### 3.3 WNT^Act^ medium significantly enhances long-term propagation of PDOs

To assess propagation capacity of PDOs in the different culture conditions, paired organoid lines from 15 patients were cultured in parallel in WNT^Act^ and WNT^Inh^ media using identical split ratios (1:2) during passages. In WNT^Inh^ medium, 12 of 15 lines degenerated or showed stagnant growth between passages 3 to 10, the remaining 3 were censored from survival analysis due to their use in other experiments. In contrast, only one of the 15 lines cultured in WNT^Act^ medium degenerated by passage 10, and the remaining 14 continued to proliferate robustly up to passage 12, at which point they were cryopreserved. Kaplan-Meier analysis confirmed the significantly improved long-term propagation (passage >10) in WNT^Act^ compared to WNT^Inh^ medium (p<0·0001, log-rank test) [**Figure 3A**, **Supplemental Table S7]**. At termination of the experiment, all PDOs generated in WNT^Act^ medium retained stable organoid morphology [**Figure 3B**]. Growth rates varied across patient-derived lines. While several PDOs (n349, n369, x357, m502, x315, m386, x365) exhibited comparable early-passage growth in both media, others proliferated faster (n319, n327 and m316) or slower (n362, n344) in WNT^Inh^ compared to WNT^Act^ media. These differences possibly reflect intrinsic patient-specific biological variability [**Figure 3C**].

**Figure 3:**
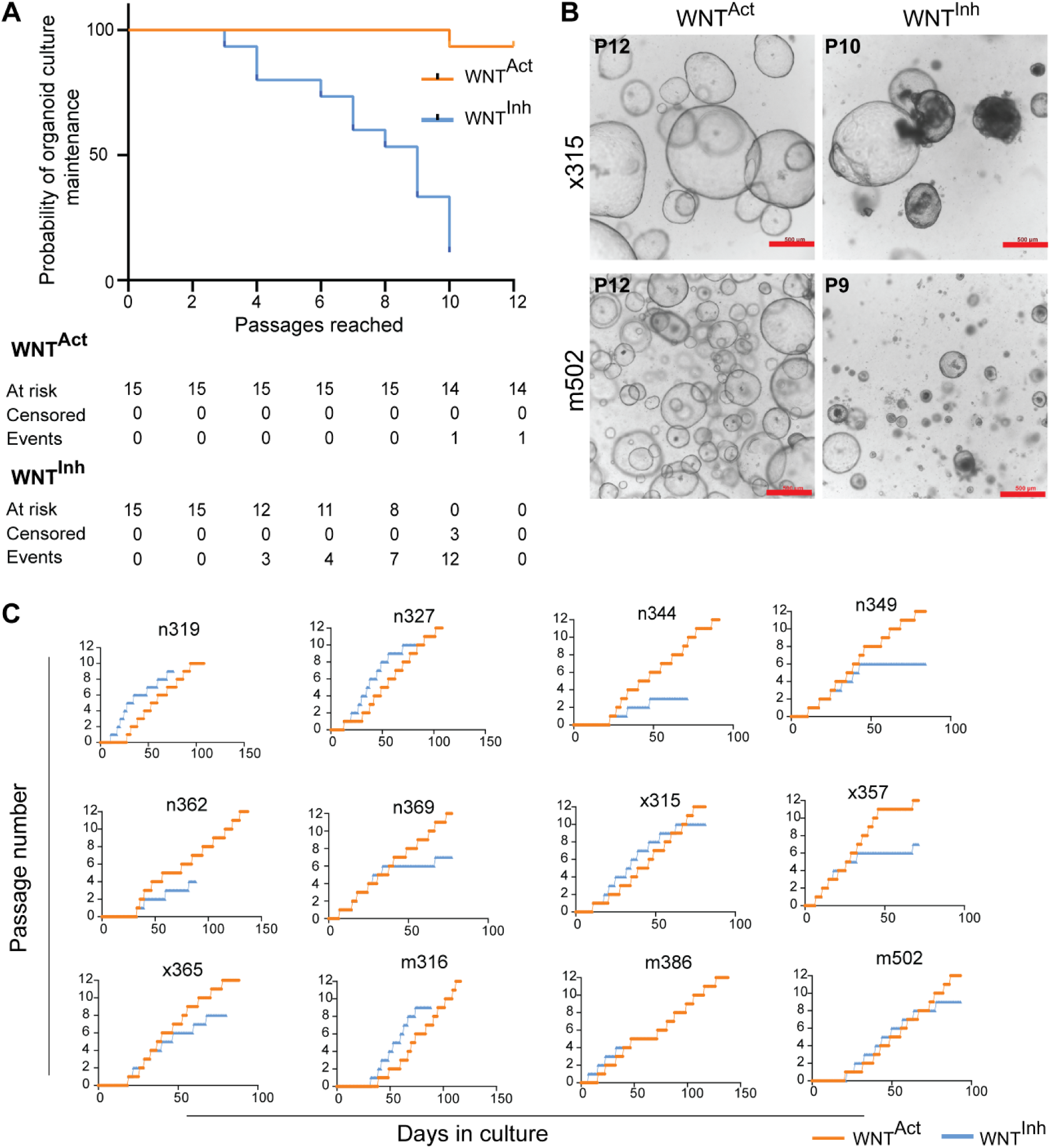
WNT^Act^ medium supports superior long-term propagation of PDOs. **(A)** Kaplan-Meier analysis of organoid propagation showing significantly improved survival of PDO lines (n = 15) cultured in WNT^Act^ medium compared with WNT^Inh^ medium p <0.0001 (log rank test). **(B)** Representative morphology of PDOs propagated in WNT^Act^ (left panel) and WNT^Inh^ (right panel) growth media from non-malignant – x315, XGC (top panel) and malignant – m502, AdCa (bottom panel) gallbladder pathologies. PDOs developed in WNT^Inh^ shows signs of degeneration and stagnant growth whereas PDOs in WNT^Act^ media were healthy and growing when cryopreserved at passage 12. **(C)** Growth curves of representative PDO lines developed in WNT^Act^ (orange) and WNT^Inh^ (blue) growth media illustrating patient-specific variability in proliferation across the two media conditions. Scale bar: 500µm. Microscope – Nikon TS2, 4X objective.**=** AdCa, adenocarcinoma; XGC or x, Xanthogranulomatous cholecystitis; PDOs, patient-derived organoids; n, non-malignant; m, malignant

### 3.4 Canonical Wnt signaling is activated in PDOs cultured in WNT^Act^ medium

To investigate transcriptional differences associated with culture conditions, mRNA-sequencing data from PDOs were subjected to principal component analysis (PCA). PDO samples clustered primarily according to tissue pathology, mirroring clustering patterns observed in the primary tissues but with clearer segregation. Organoids derived from invasive malignant tissues (MAL) segregated from non-malignant ones (non-MAL), whereas XGC organoids frequently clustered near MAL and pre-invasive neoplasm (PIN) samples. A secondary clustering effect in PDOs was attributable to the culture medium [**Figure 4A**], consistent with previous observations (10). Gene set enrichment analysis (GSEA) using an 18 gene canonical Wnt target panel [**Supplemental Table S5**] demonstrated significant enrichment of Wnt target gene expression in PDOs cultured in WNT^Act^ compared to the WNT^Inh^ (NES = 1.5, FDR = 0.01) ones [**Figure 4B**]. qRT-PCR confirmed significant upregulation of canonical Wnt targets *LGR5, TCF7, AXIN2, LEF1,* and *MMP7* in WNT^Act^ but not in WNT^Inh^ cultures [**Figure 4C**]. Expression of pluripotency-associated stem cell markers, *NANOG, POU5F1* and *SOX2* were not be detected in the developed PDO lines suggesting Wnt activation did not induce a prominent stem-like transcriptional state [Data not shown]. Together, these data confirm activation of the canonical Wnt pathway in PDOs cultured in WNT^Act^ medium but the organoid gene expression is primarily determined by the source tissue pathology.

**Figure 4:**
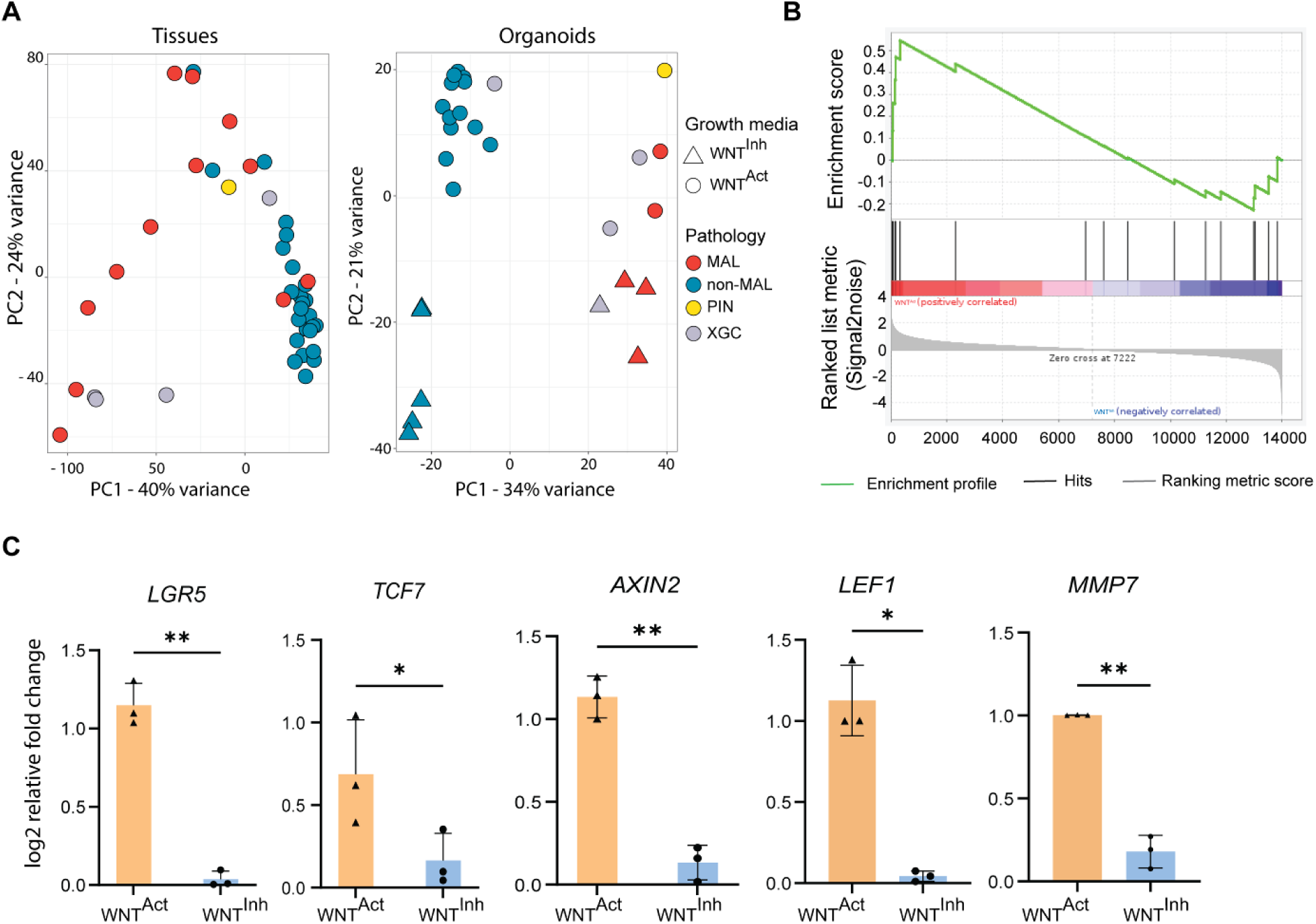
Canonical Wnt signaling is activated in WNT^Act^ cultured PDOs. **(A)** Principal component analysis of mRNA sequencing data from gallbladder PDOs (right) and primary tissue samples (left) based on the gene expression of top 1000 highly variable genes, showing clustering primarily determined by source tissue pathology. Pathology groups have been indicated by the colours and growth media types by the shapes. **(B)** Gene set enrichment analysis demonstrating enrichment of the canonical Wnt target genes in PDOs developed in WNT^Act^ media. **(C)** qRT-PCR validating of canonical Wnt targets *LGR5, TCF7, AXIN2, LEF1* and *MMP7* in GCOs developed in WNT^Act^ media, plotted Mean +/- SD (n = 3 samples), * (p < 0.05), ** (p < 0.01); paired t-test. GSEA, gene set enrichment analysis; GCOs, gallbladder cholangiocyte organoids; PDOs, patient-derived organoids, MAL, malignant group; non-MAL, non-malignant group; PIN, pre-invasive neoplasm; XGC, xanthogranulomatous cholecystitis.

### 3.5 WNT^Inh^ grown GBCO lines more consistently retain dysplastic histological features

To evaluate histological fidelity, paired PDO cultures derived from the same patient tissue but grown in WNT^Act^ and WNT^Inh^ media, were analyzed by hematoxylin–eosin–staining to assess retention of cytological and architectural features of the original tissue. PDOs derived from non-malignant tissues (CC, XGC, normal gallbladder) exhibited single-layered epithelial organization with central lumens and normal nucleus-to-cytoplasm (N:C) ratios, under both culture conditions [**Supplemental Figure S1A, Supplemental Table S8**]. In contrast, PDOs derived from pre-invasive or invasive lesions demonstrated differential retention of dysplastic features depending on which medium was used. All 12 GBCO lines cultured in WNT^Inh^ medium retained features of moderate-to-high dysplasia, consistent with their source tissues [**Figure 5A–B, Supplemental Figures S1B–S1C, Supplemental Table S8**]. However, only 6 (p379, m239, m289, m290, m385, m502) of the 12 paired lines cultured in WNT^Act^ medium retained comparable dysplastic characteristics. The remaining lines displayed reduced or absent dysplastic features [**Figure 5B, Supplemental Figure S1C, Table S8**]. The evaluated dysplastic features included epithelial multilayering, lumen filling, nuclear pleomorphism, hyperchromasia, chromatin clumping, prominent nucleoli, increased N:C ratio, and mitotic activity.

**Figure 5:**
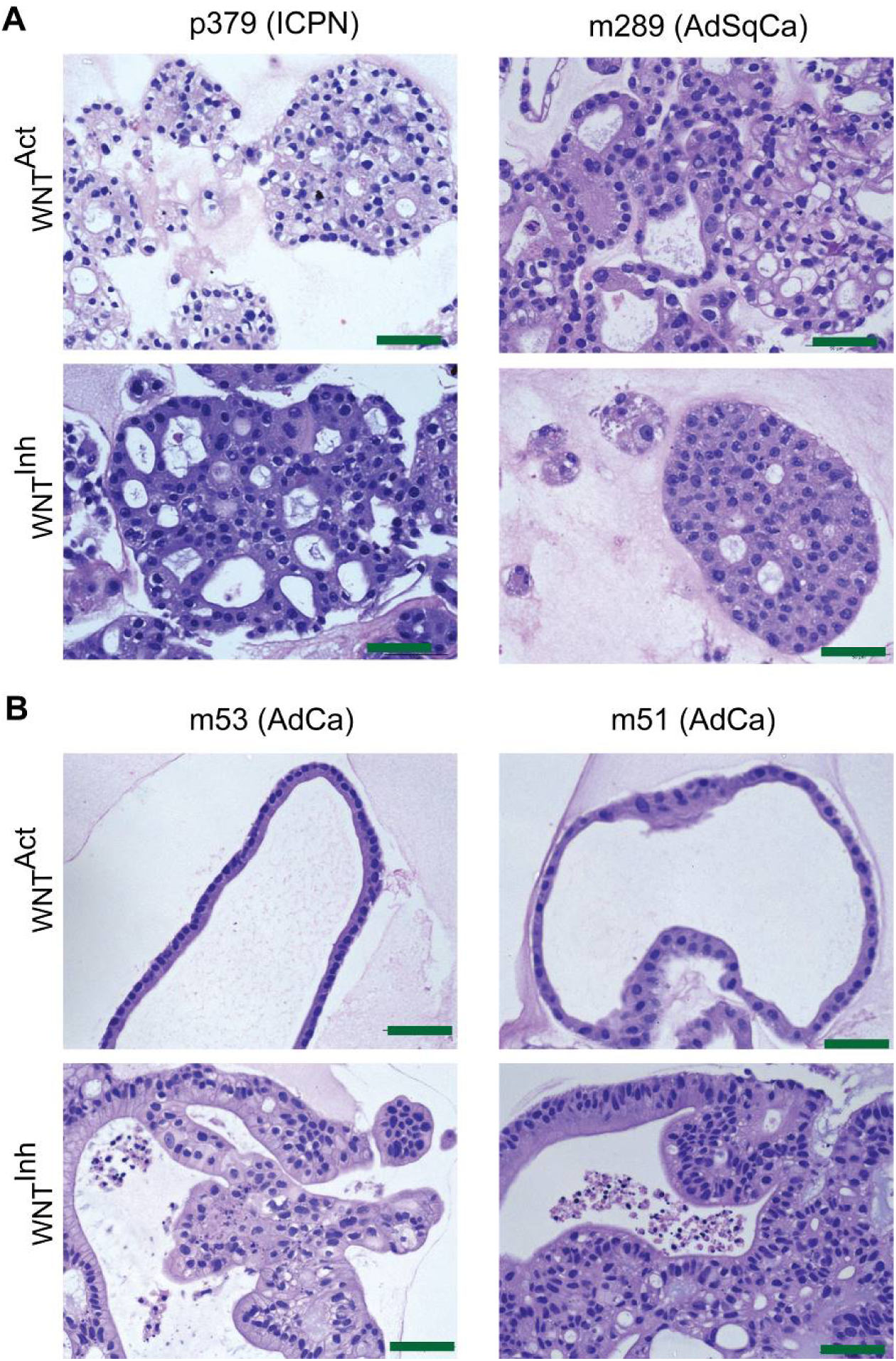
Differential retention of dysplastic histology in GBCO lines developed in WNT^Inh^ and WNT^Act^ growth medium. Representative hematoxylin-eosin stained FFPE sections of GBCOs. **(A)** Representative images of stained GBCO lines that retained cytological and architectural dysplastic features both in WNT^Act^ and WNT^Inh^ media **(B)** Representative images of stained GBCO lines that retained dysplastic features only in WNT^Inh^ medium but not in WNT^Act^ medium. Scale bar - 50μm. Microscope – Leica LMD7, 40X objective. AdCa, adenocarcinoma; ICPN, intra-cholecystic papillary tubular neoplasm; FFPE, formalin fixed paraffin embedded; HE, hematoxylin eosin; GBCOs, gallbladder carcinoma organoids

### 3.6 WNT^Act^ culture enhances gemcitabine sensitivity in GBCO lines

To investigate whether Wnt modulation influences therapeutic response, paired GBCO lines retaining dysplastic morphology were treated with gemcitabine, a standard of care chemotherapeutic agent for GBC. In all three tested pairs (two AdCa: m290, m502; one AdSqCa: m289), PDOs cultured in WNT^Act^ medium showed significantly greater sensitivity to gemcitabine compared to their WNT^Inh^ counterparts (p <0·.001, F-test) [**Figure 6A, Supplemental Figure S2A**]. To assess whether differential drug response could be attributed to proliferation differences, Ki-67 immunohistochemistry was performed in these three pairs of GBCOs. Ki67 expression lacked a consistent pattern between the two culture conditions [**Supplemental Figure S2B**]. Ki-67 index was higher in WNT^Act^ (∼30%) as compared to WNT^Inh^ (∼25%) in m289. In m502, Ki65 index was higher in WNT^Inh^ (∼70%) as compared to WNT^Act^ (∼ 30%). For m290, Ki67 index was similar in both lines (∼15% each). These findings suggest that proliferation rate alone cannot explain the observed differences in drug sensitivity.

**Figure 6:**
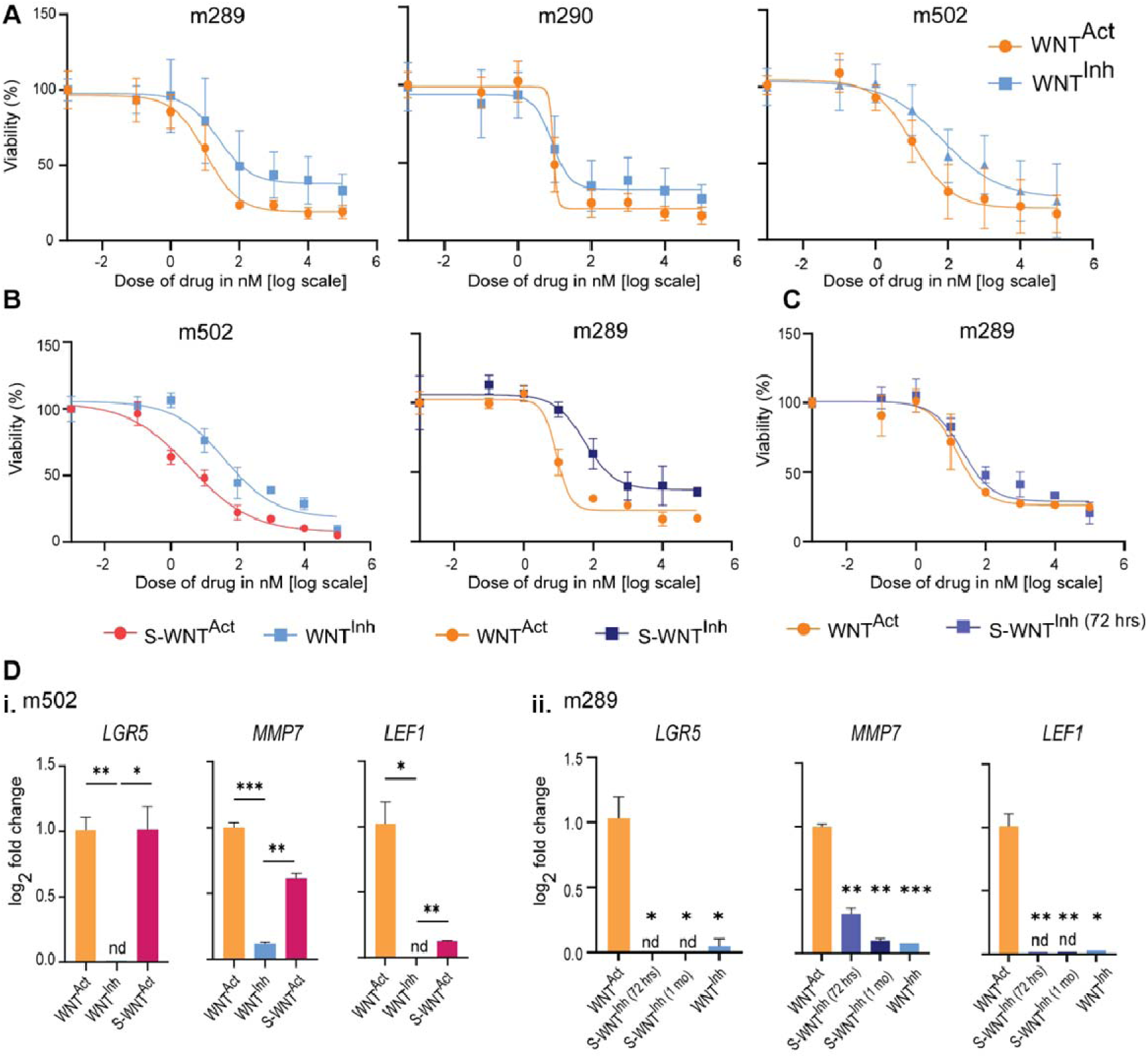
Wnt signaling modulates gemcitabine sensitivity in gallbladder cancer organoids. **(A)** Dose-response curves showing significantly increased gemcitabine sensitivity in PDOs generated in WNT^Act^ (orange) compared to WNT^Inh^ (blue) medium in three malignant gallbladder patients (m289, m290 and m502) (p<0.0001 for each pair, F-test). 8000 PDO cells were seeded in 5% Geltrex^TM^ and treated with DMSO or different concentrations of drug with 10-fold serial dilutions. Viability was assessed using fluorometric CyQUANT^TM^ Direct Cell Proliferation Assay 72 hours post-treatment. Dose response curves were generated using curve fitting to non-linear four-parameter logistic regression equation in GraphPad Prism 10. **(B)** Dose response curves following media switching experiments demonstrating reversible modulation of gemcitabine sensitivity. Left: GBCO line m502 switched into S-WNT^Act^ (> 1 month) (red) responded significantly better compared to the parent line continued to be cultured in WNT^Inh^ (blue) (p<0.0001, F-test). Right: GBCO line m289 developed and maintained in WNT^Act^ (orange) responded significantly better than when switched into S-WNT^Inh^ medium (dark blue) for >1 month (p<0.0001, F-test). **(C)** Dose response curves performed after 72 hours of switching media showing no change in sensitivity. No significant difference in response to gemcitabine was observed in PDO m289 developed and maintained in WNT^Act^ medium (orange) or when switched into S-WNT^Inh^ medium (blue) for 72 hours before the assay. (p>0.05, F-test). **(D)** qRT-PCR analysis of canonical Wnt targets: *LGR5, LEF1* and *MMP7* following media switching. **(i)** Upregulation of the targets in GBCO line m502 when grown in WNT^Act^ or S-WNT^Act^ media compared to its WNT^Inh^ counterpart (>1 month). **(ii)** Downregulation of the targets in GBCO line m289 when grown in WNT^Act^ compared to WNT^Inh^ medium and when switched to S- WNT^Inh^ for 72-hours or 1-month. Mean +/- SEM, * (p<0.05), ** (p<0.01), ***(p<0.001); paired t-test. Reference gene used is *PPIA*. GBCO, gallbladder carcinoma organoids; DMSO, dimethyl sulfoxide; nd, not detected; S-WNT^Act^, growth medium switched from WNT^Inh^ to WNT^Act^; S-WNT^Inh^, growth medium switched from WNT^Act^ to WNT^Inh^

To determine whether the Wnt-dependent drug response was reversible, media-switch experiments were performed. A GBCO line (m502) initially cultured for 80 days (six passages) in WNT^Inh^ medium was switched into WNT^Act^ medium (S-WNT^Act^) and cultured for more than one month prior to drug response testing. S-WNT^Act^ had significantly increased gemcitabine sensitivity compared to the passage-matched WNT^Inh^ control [**Figure 6B-left, Supplemental Figure S3A**]. Conversely, another GBCO line (m289) initially cultured in WNT^Act^ medium for 78 days (eight passages) was switched into WNT^Inh^ medium (S-WNT^Inh^) and cultured for more than 1 month in S-WNT^Inh^ before drug response testing. This resulted in decreased gemcitabine sensitivity in S-WNT^Inh^ relative to the passage matched WNT^Act^ control **[Figure 6B-right, Supplemental Figure S3B]**. Importantly, organoids maintained in switched media retained high-grade dysplastic features at the time of drug testing **[Supplemental Figure S3C]**. Consistent with earlier observations, Ki-67 staining did not correlate with changes in drug response **[Supplemental Figure S3D]**. No alterations in drug response was observed when drug response assay was performed 72 hours after switching media for m289 suggesting phenotypic changes require prolonged culture adaptation. [**Figure 6C].** However, qRT-PCR showed significant changes in canonical Wnt target gene expression (*LGR5*, *MMP7* and *LEF1)* following media switching, detectable at 72 hours **[Figure 6D]**. Together, these results suggest that Wnt-mediated modulation of drug response arises from delayed cellular adaptation rather than immediate pathway activation or direct interference from presence/absence of media components.

## Discussion

In this study we systematically investigated how modulation of canonical Wnt signaling influences the establishment, phenotype and therapeutic responsiveness of gallbladder PDO. Using a large biorepository of gallbladder PDOs, representing a broad spectrum of benign and malignant gallbladder pathologies (10), we directly compared organoids grown under two previously reported culture conditions, with activated (7) and inhibited (9) canonical Wnt. Gallbladder PDOs were generated from malignant and non-malignant tissues with comparable efficiency in both culture conditions. Organoid morphology was largely determined by the pathology of the source tissue rather than the culture medium, consistent with previous studies in other epithelial organoid systems demonstrating that PDOs retain intrinsic tissue-specific characteristics (11). Functional assays confirmed that organoids maintained key features of biliary epithelial cells, including active transporter function and expression of canonical cholangiocyte markers. These observations indicate that both media support the establishment of gallbladder epithelial organoids while preserving fundamental aspects of biliary epithelial identity.

Despite comparable initial establishment efficiency, we observed a striking difference in the long-term propagation potential of PDOs cultured under the two conditions. WNT^Act^ medium supported robust long-term expansion of most organoid lines, whereas cultures maintained in WNT^Inh^ medium frequently displayed stagnant growth or degeneration at relatively early passages. Transcriptomic analyses confirmed activation of canonical Wnt signaling in WNT^Act^-grown PDOs, including enrichment of established Wnt target genes such as *LGR5*, *TCF7*, *AXIN2*, and *LEF1*. These findings are consistent with the well-established role of Wnt signaling in maintaining epithelial progenitor cell proliferation and organoid growth across multiple tissues, including intestinal, hepatic, and biliary organoid models (12,13,6). In this study, activation of canonical Wnt signaling did not induce detectable expression of pluripotency-associated stem cell markers, suggesting that Wnt activation in this context promoted proliferative capacity without inducing a broadly dedifferentiated stem-like state. While WNT^Act^ conditions enhanced proliferative capacity, WNT^Inh^ cultures more consistently preserved dysplastic histological features in organoids derived from malignant tissues. All evaluated gallbladder cancer organoid (GBCO) lines cultured in WNT^Inh^ medium retained moderate-to-high-grade dysplasia closely resembling the corresponding source tissues, whereas only half of the paired lines maintained comparable dysplastic characteristics in WNT^Act^ medium. These findings suggest that high levels of Wnt signaling may partially remodel epithelial architecture or promote expansion of less dysplastic cellular populations within heterogeneous tumor-derived cultures. Similar observations have been reported in other organoid systems in which growth factor–rich media promoted selective expansion of specific epithelial subpopulations (14). Thus, a trade-off appears to exist between optimal conditions for long-term expansion and faithful preservation of tumor histopathology.

A key finding was that Wnt pathway modulation significantly influenced the ex vivo therapeutic response. PDOs cultured in WNT^Act^ medium exhibited greater sensitivity to gemcitabine compared with their WNT^Inh^ counterparts. This difference in drug response was reversible when cultures were switched between media conditions, demonstrating that the phenotype is not permanent rather reflects dynamic cellular adaptation to the signaling environment. Media-switch experiments further revealed that although Wnt target gene expression changed rapidly following pathway modulation, alterations in gemcitabine sensitivity required a longer adaptation period. This temporal separation suggests that drug response modulation may arise from indirect downstream transcriptional or metabolic reprogramming rather than from immediate signaling effects. The lack of correlation between Ki-67 proliferation indices and gemcitabine sensitivity indicates that differential drug response is unlikely to be explained solely by differences in cell proliferation rates. Instead, Wnt signaling may influence additional cellular processes relevant to chemotherapeutic susceptibility. Further mechanistic studies are required to delineate the molecular pathways linking Wnt signaling activity to chemotherapy sensitivity in gallbladder cancer cells.

This study has several limitations. First, although the PDO biobank includes a large number of lines overall, functional drug response analyses were performed in a limited number of paired organoid models. Expanding these experiments to a larger cohort will be important to determine the generalizability of the observed Wnt-dependent drug sensitivity. Second, organoid cultures lack components of the tumor microenvironment, including immune and stromal cells, which may influence drug response in vivo. Finally, while transcriptomic analyses confirmed activation of canonical Wnt signaling in WNT^Act^ cultures, the precise downstream mechanisms responsible for altered histological features and drug response remain to be elucidated. Despite these limitations, our findings have important implications for the use of gallbladder PDOs in translational research. This study demonstrates that canonical Wnt signaling exerts profound effects on long-term organoid propagation, phenotypic fidelity, and chemotherapy response. Practically, WNT^Inh^ conditions are optimal for preserving dysplastic and architectural features relevant to disease modeling, whereas WNT^Act^ conditions are superior for long-term propagation. To achieve long term propagation of gallbladder cancer PDO lines without compromising in either proliferation or preservation of disease features, a judiciously balanced hybrid culture approach can be tested. Culture establishment and initial maintenance can be under WNT^Inh^ to preserve disease features more consistently while intermittent expansion under WNT^Act^ for brief period may be helpful to prevent decline in proliferation. To choose suitable culture condition between WNT^Act^ and WNT^Inh^ for therapeutic response prediction, a comparative clinical response correlation analysis has to be performed. Finally, the large PDO biobank established in this study provides a valuable resource for future investigations of gallbladder cancer biology and therapeutic testing. Overall, this work provides a comprehensive comparative framework for Wnt pathway modulation in gallbladder organoid cultures and underscores the importance of tailoring culture conditions to the scientific or clinical question at hand. By illuminating the interplay between signaling environment, cellular phenotype, and therapeutic vulnerability, these findings will help guide the design and interpretation of future studies using gallbladder PDO models.

## Resource Availability

### Materials availability

No new reagents have been generated in this study. Banked organoid lines are available on request to the corresponding author against a complete material transfer agreement and reasonable compensation for processing and shipping.

### Data and code availability

No original code has been developed in the current study. mRNAseq data is available at NCBI SRA (PRJNA1209630).

## Supporting information

Supplemental Figures with Legends

## Acknowledgement

This work is part funded by a Wellcome-DBT India Alliance Margdarshi Fellowship (IA/M/12/500755) to VS and a Tata Consultancy Services Foundation and Tata Trust core grant to TTCRC and Ministry of Education, India PhD Fellowship to AkD (Ankita Dutta). We thank Cancer Genomics Laboratory, IT unit, Clinical Research Unit, Biobank (TiMBR), admin and house-keeping teams at TTCRC. We appreciate Pathology, GI-HPB, Operation Theatre teams and nurses at TMC for their support. We express gratitude to the patients and family for donating clinical samples.

## Author contribution

Conceptualization: VS, AD (Anindita Dutta), DGS; Experiment design: VS, AD, DGS; Performing experiments and data collection: AkD, PG, NC, AkS; Coordination, patient consenting and clinical sample collection: PB, SSG, MKR, SB; Data analysis and interpretation: AkD, PG, AVS, DGS, US, RS, PR, VS; Supervision: VS, DGS, AD, RS, US, DG, MKR, SB; Writing original draft: AkD, DGS; Review and critical editing: DGS, VS; Funding acquisition: VS;

## References

1. Sepe LP, Hartl K, Iftekhar A, Berger H, Kumar N, Goosmann C, et al. Genotoxic Effect of Salmonella Paratyphi A Infection on Human Primary Gallbladder Cells. mBio. 2020 Sep 22;11(5):e01911–20. doi:10.1128/mBio.01911-20

2. Saito Y, Muramatsu T, Kanai Y, Ojima H, Sukeda A, Hiraoka N, et al. Establishment of Patient-Derived Organoids and Drug Screening for Biliary Tract Carcinoma. Cell Rep. 2019 Apr 23;27(4):1265–1276.e4. doi:10.1016/j.celrep.2019.03.088

3. Clevers H. Modeling Development and Disease with Organoids. Cell. 2016 Jun 16;165(7):1586–97. doi:10.1016/j.cell.2016.05.082

4. Nusse R, Clevers H. Wnt/β-Catenin Signaling, Disease, and Emerging Therapeutic Modalities. Cell. 2017 Jun 1;169(6):985–99. doi:10.1016/j.cell.2017.05.016 PubMed PMID: 28575679.

5. Dutta A, Chowdhury N, Chandra S, Guha P, Saha V, GuhaSarkar D. Gallbladder cholangiocyte organoids. Biol Cell. 2025 Feb;117(2):e2400132. doi:10.1111/boc.202400132

6. Rimland CA, Tilson SG, Morell CM, Tomaz RA, Lu WY, Adams SE, et al. Regional Differences in Human Biliary Tissues and Corresponding In Vitro-Derived Organoids. Hepatology. 2021 Jan;73(1):247–67. doi:10.1002/hep.31252

7. Yuan B, Zhao X, Wang X, Liu E, Liu C, Zong Y, et al. Patient-derived organoids for personalized gallbladder cancer modelling and drug screening. Clin Transl Med. 2022 Jan;12(1):e678. doi:10.1002/ctm2.678

8. Sampaziotis F, Justin AW, Tysoe OC, Sawiak S, Godfrey EM, Upponi SS, et al. Reconstruction of the mouse extrahepatic biliary tree using primary human extrahepatic cholangiocyte organoids. Nat Med. 2017 Aug;23(8):954–63. doi:10.1038/nm.4360

9. Tysoe OC, Justin AW, Brevini T, Chen SE, Mahbubani KT, Frank AK, et al. Isolation and propagation of primary human cholangiocyte organoids for the generation of bioengineered biliary tissue. Nat Protoc. 2019 Jun;14(6):1884–925. doi:10.1038/s41596-019-0168-0

10. Dutta A, Chowdhury N, Selvarajan AV, Guha P, Banerjee P, Kar A, et al. An Annotated Living Organoid Biobank for Studying Gallbladder Diseases and Drug Responses. Dig Dis Sci. 2026 Feb 5. doi:10.1007/s10620-026-09707-x

11. Dekkers JF, van Vliet EJ, Sachs N, Rosenbluth JM, Kopper O, Rebel HG, et al. Long-term culture, genetic manipulation and xenotransplantation of human normal and breast cancer organoids. Nat Protoc. 2021 Apr;16(4):1936–65. doi:10.1038/s41596-020-00474-1

12. Sato T, Vries RG, Snippert HJ, van de Wetering M, Barker N, Stange DE, et al. Single Lgr5 stem cells build crypt-villus structures in vitro without a mesenchymal niche. Nature. 2009 May 14;459(7244):262–5. doi:10.1038/nature07935

13. Huch M, Gehart H, van Boxtel R, Hamer K, Blokzijl F, Verstegen MMA, et al. Long-term culture of genome-stable bipotent stem cells from adult human liver. Cell. 2015 Jan 15;160(1–2):299–312. doi:10.1016/j.cell.2014.11.050

14. Usman OH, Zhang L, Xie G, Kocher HM, Hwang CI, Wang YJ, et al. Genomic heterogeneity in pancreatic cancer organoids and its stability with culture. NPJ Genomic Med. 2022 Dec 19;7(1):71. doi:10.1038/s41525-022-00342-9

