## Supplemental Figures with Legends for "Wnt stimulation and inhibition in the development and phenotype of patient-derived gallbladder organoids"

**Supplemental Figure S1**

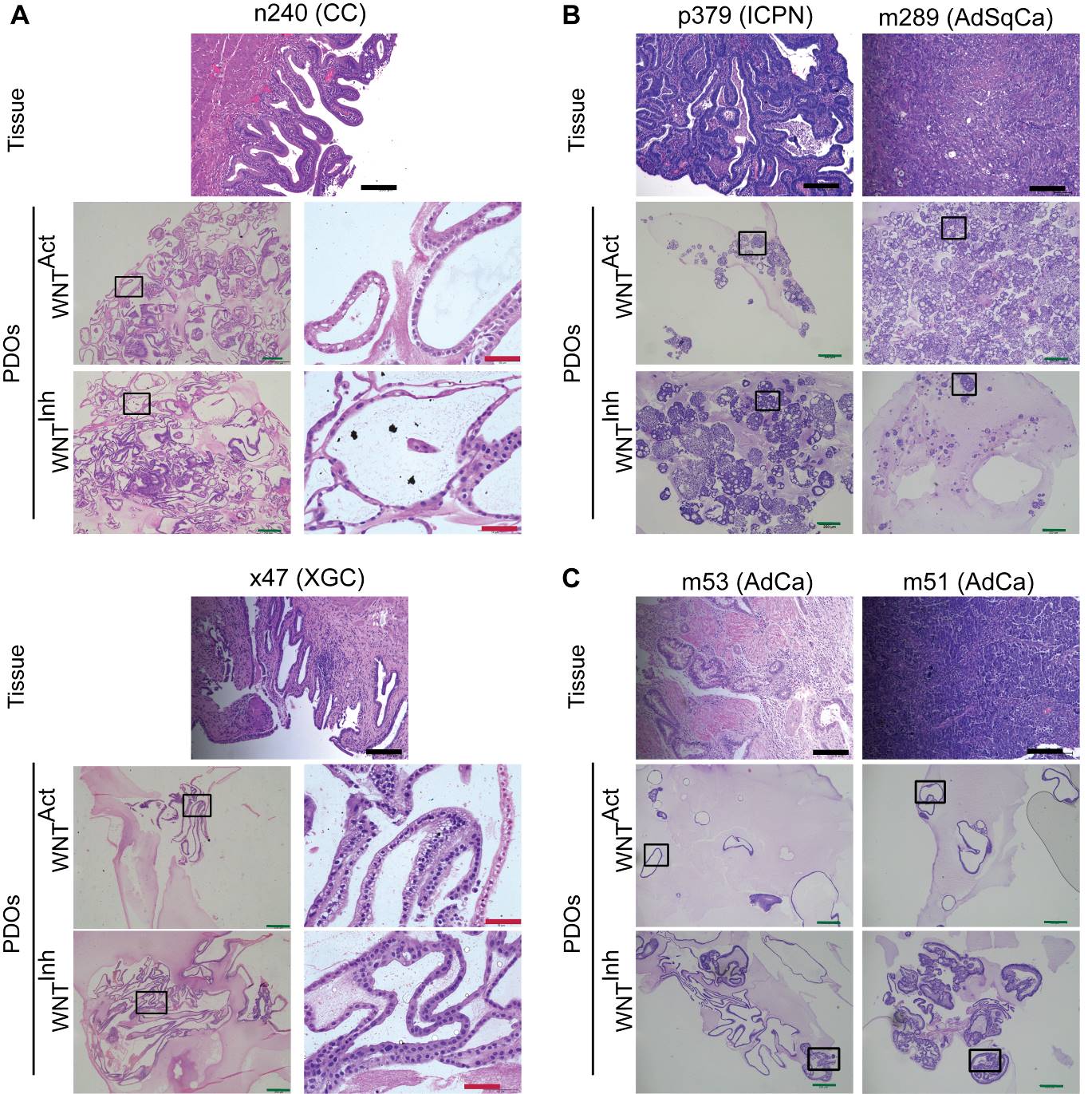

**Supplemental Figure S1: - GBCOs developed in** **WNT^Inh^ medium retain the histopathological features of the source tissue more consistently compared to the paired WNT^Act^ grown lines**

Representative bright field images of the HE-stained FFPE sections of the tissues and corresponding PDOs developed in WNT^Act^ and WNT^Inh^ media as indicated. The images show **(A)** GCOs grown from non-malignant pathologies, where there were no cytological or architectural differences observed between the paired organoids grown in two different media, **(B)** GBCOs retained cytological and architectural features of dysplasia in both the culture media and **(C)** GBCOs that retained the dysplasia features only in WNT^Inh^ but not in WNT^Act^. Higher magnification images of the marked areas of PDOs indicated in A are shown in the corresponding right panels. The higher magnification images for B and C are shown in Figure 5. Microscope – Leica LMD7, 5X and 10X objective. Scale bar - 100μm (black), 250μm (red).

CC, chronic cholecystitis; XGC, Xanthogranulomatous cholecystitis; PDOs; patient derived organoids. Rest all abbreviations used here are discussed in Figure 5

**Supplemental Figure S2**

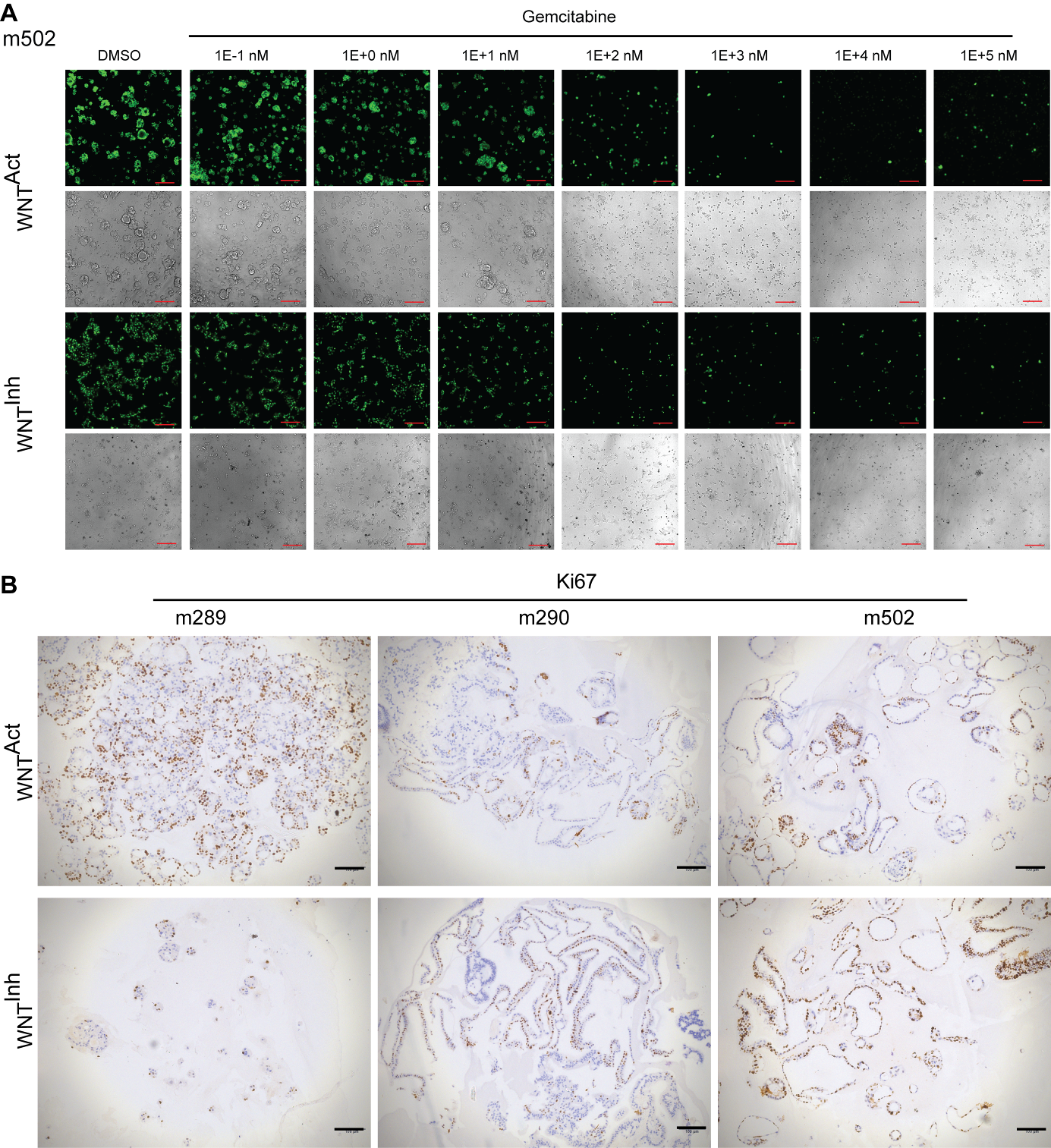

**Supplemental Figure S2: PDOs developed in WNT^Act^ growth medium responded better to gemcitabine compared to WNT^Inh^ grown lines irrespective of their comparative proliferation rate**

**(A)** Fluorescence (top row) and bright-field (bottom row) images of DMSO or gemcitabine treated GBCOs (m502, adenocarcinoma) developed in WNT^Act^ and WNT^Inh^ growth media at 72-hours post treatment. Cells were stained with CyQUANT^TM^ dye prior to imaging. Treatment agents and drug doses are indicated above each respective image. **(B)** Immunohistochemical analysis showing the Ki67 expression in the GBCOs cultured in WNT^Act^ (top) and WNT^Inh^ (bottom) media for m289 (left), m290 (middle) and m502 (right). Proliferation status as measured by Ki67 indices lacked correlation with drug response. Scale bar - 200 µm (red), 100 µm (black). Microscope - ImageXpress Micro Confocal High-Content Imaging System (Molecular Devices), wide-field mode, 10X objective (A); Leica LMD7, 10X objective (B).

GBCO, gallbladder carcinoma organoids; DMSO, dimethyl sulfoxide

**Supplemental Figure S3**

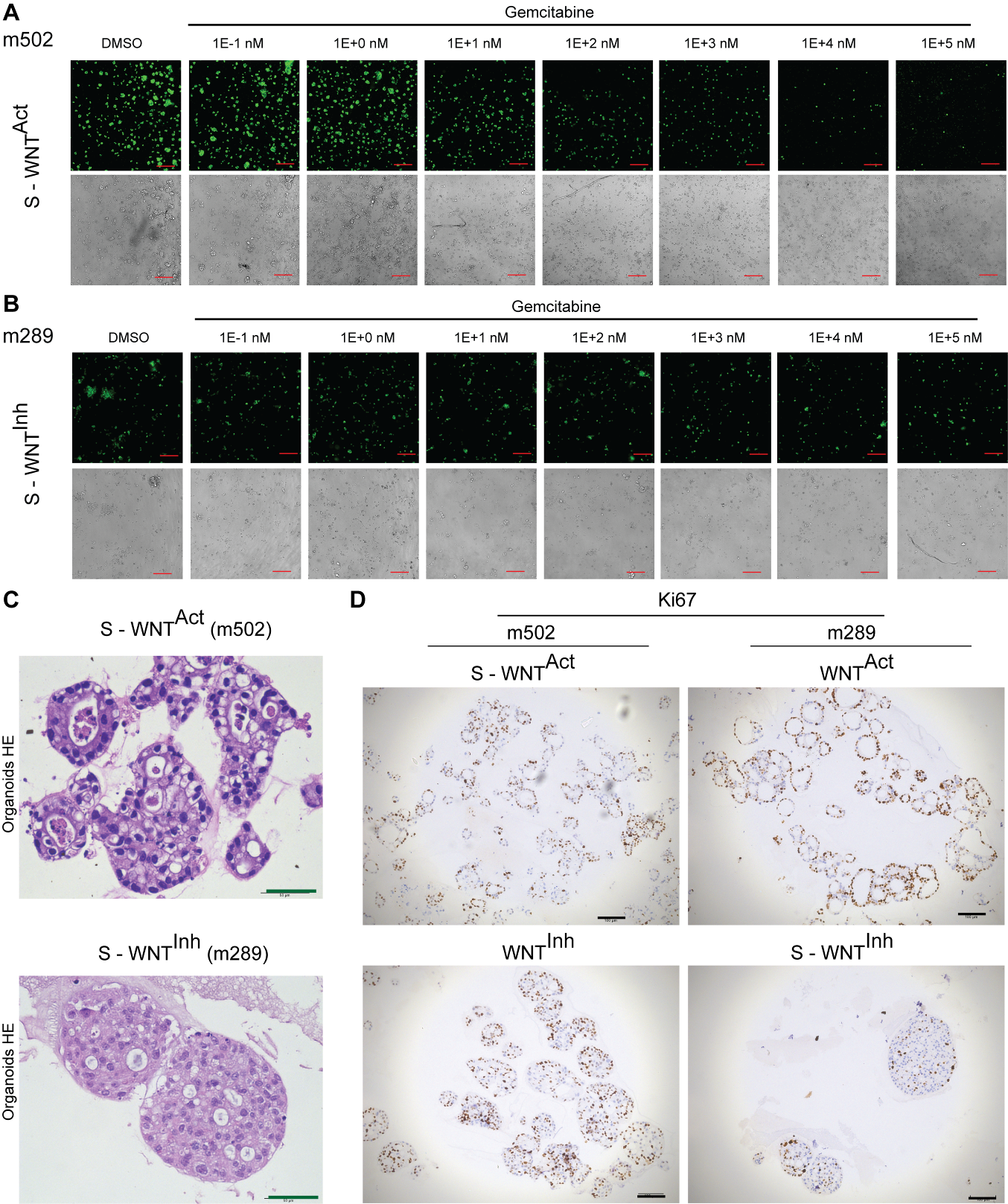

**Supplemental Figure S3: PDOs developed in switched growth media showed altered drug response phenotype irrespective of the proliferation status**

Fluorescence (top row) and bright-field (bottom row) images of DMSO or gemcitabine treated GBCOs developed in **(A)** S-WNT^Act^ from patient m502 and **(B)** S–WNT^Inh^ from patient m289 at 72-hours post treatment. Cells were stained with CyQUANT^TM^ dye prior to imaging. Treatment agents and drug doses are indicated above each respective image. **(C)** Representative images of HE-stained FFPE sections of the GBCOs from m502 cultured in S-WNT^Act^ medium (top panel) and m289 cultured in S–WNT^Inh^ medium (bottom panel) retaining cytological and architectural features of dysplasia. **(D)** Immunohistochemical analysis showing the Ki67 expression in the GBCOs cultured in S-WNT^Act^ and WNT^Inh^ media for m502 (left) and S–WNT^Inh^ and WNT^Act^ media for m289 (right). Proliferation status lacked correlation with drug response. Scale bar - 200 µm (red), 100 µm (black) and 50 µm (green). Microscope - ImageXpress Micro Confocal High-Content Imaging System (Molecular Devices), wide-field mode, 10X objective (A and B); Leica LMD7, 10X (C) and 40X (D) objective.

S-WNT^Act^, growth medium switched from WNT^Inh^ to WNT^Act^; S-WNT^Inh^, growth medium switched from WNT^Act^ to WNT^Inh^
